# Phased nanopore assembly with Shasta and modular graph phasing with GFAse

**DOI:** 10.1101/2023.02.21.529152

**Authors:** Ryan Lorig-Roach, Melissa Meredith, Jean Monlong, Miten Jain, Hugh Olsen, Brandy McNulty, David Porubsky, Tessa Montague, Julian Lucas, Chris Condon, Jordan Eizenga, Sissel Juul, Sean McKenzie, Sara E. Simmonds, Jimin Park, Mobin Asri, Sergey Koren, Evan Eichler, Richard Axel, Bruce Martin, Paolo Carnevali, Karen Miga, Benedict Paten

## Abstract

As a step towards simplifying and reducing the cost of haplotype resolved *de novo* assembly, we describe new methods for accurately phasing nanopore data with the Shasta genome assembler and a modular tool for extending phasing to the chromosome scale called GFAse. We test using new variants of Oxford Nanopore Technologies’ (ONT) PromethION sequencing, including those using proximity ligation and show that newer, higher accuracy ONT reads substantially improve assembly quality.

## Introduction

Phasing of genome assemblies enables a variety of clinically motivated analyses and population studies. For clinical and biological studies, there are many transcriptional and translational outcomes that could result from a given set of mutations in the same gene or regulon, depending on whether they are actually co-occurring on the same molecule of DNA^1–3^. Additionally, understanding how loci are linked enables imputation, and therefore a high quality set of phased variants can serve as a catalyst for much larger volume experiments, creating greater statistical power for disease association^4,5^. Population genetics also benefits from haplotype information because it provides a means to estimate recombination and gene flow^6^. As a consequence of its many applications, a significant portion of recent efforts in human genomics have become focused on generating a high quality, genome-wide set of phased variants^7,8^.

Methods for read-based phasing generally consist of two approaches that either rely on an existing assembly or use reads directly for creating a phased consensus. When read length or accuracy is limited, mapping-based methods are essential, because mapping is required to find a set of candidate sites that share reads. After mapping, reads are usually phased by finding a partition which maximizes the consistency of shared reads among the alleles^9–11^. Trio binning is sometimes feasible prior to assembly, which simplifies the problem by labeling reads directly with exact subsequences (k-mers) from parental data, but its applications have been limited by repetitive or homozygous regions which have few haplotype-specific k-mers^12^.

In contrast to reference based phasing, *de novo* methods for read-based phasing generate candidate variants internal to the assembler, following read overlap^13,14^. The advantage of assembly based methods is that they do not fall victim to the problem of reference bias, and can therefore identify variants that would otherwise map poorly, as with repetitive regions or large duplications and inversions^15,16^. Alignment-based methods can work around this issue by using a draft assembly instead of an existing reference.

Since the basis of assembly phasing lies in the overlapping of reads, highly repetitive or homozygous regions limit phasing in Shasta and other assemblers such as Hifiasm and Verkko^13,14^. To address this, proximity ligation data types have been used for extending phasing beyond the length of a typical read, without the need for parental sequencing^17^. Data types like Hi-C and Pore-C exploit the physical packing of chromatin in the nucleus to ligate proximal regions of DNA molecules, which can be hundreds of millions of base pairs apart, linearly^18,19^. In this paper, we take an approach to proximity ligation based phasing methods, which leverages assembly graph information to phase and extend partial haplotypes.

To date, there is only one published example of a nanopore assembler which claims to produce phased haplotypes^20^, and a paper prior to that concluded that nanopore *de novo* phasing is not practical ^21^. However, the Shasta assembler has kept pace with recent advances in nanopore sequencing by iterating through multiple approaches for phasing, and we now present a new version which uses long reads to phase variation observed in its sequence graph. Using nanopore sequencing for both long reads and proximity ligated reads, we demonstrate a logistically simpler alternative to hybrid and HiFi based approaches, while attaining comparable switch and hamming rates (Figure 1). We test the limits of cost efficiency using a single flowcell of PromethION R10, and attempt to maximize assembly continuity with high coverage ultra-long reads.

**Figure 1:**
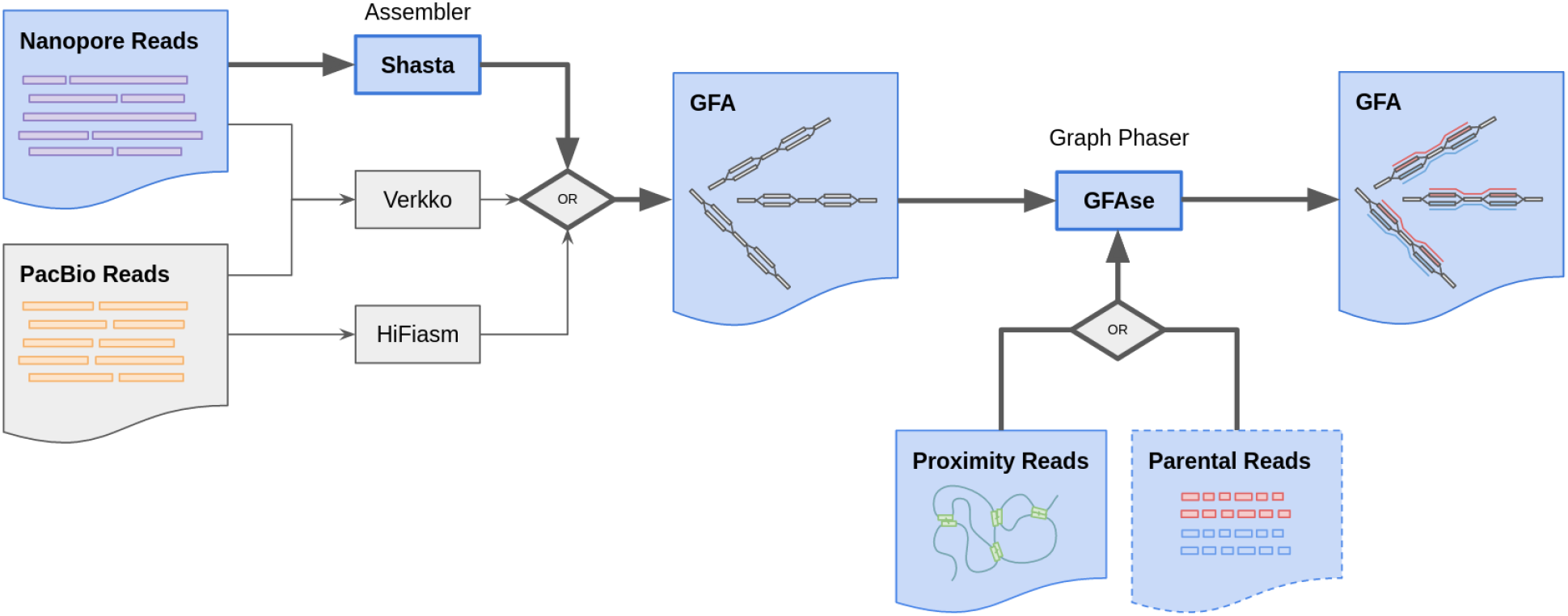
Summary of *de novo* phasing pipeline using Shasta and GFAse. Shasta performs *de novo* assembly and phases to the extent that is supported by informative variants in the nanopore reads. GFAse then takes a partially phased assembly GFA and extends phasing using orthogonal phasing information. GFAse can perform phasing based on any alignable data type (HiC, Pore-C, etc.). For Shasta graphs, GFAse can also use parental sequencing. The pathways with bolded arrows and blue fill are the methods that are previously undescribed.

## Results

### Long read sequencing

To address variability in read length, coverage, and accuracy, these results evaluate phasing in 6 different combinations of library preparation and chemistry. The effect of read accuracy on phasing is addressed with differing nanopore chemistries: R9 and R10. For the datasets evaluated, R9 has median accuracy of 95.7-96.1% (Figure 2a) and R10 has 98.3-98.9% median accuracy. In ONT R9, three different length distributions and coverages were assembled. The datasets have minimum read lengths of 10Kbp, 35Kbp, and 100Kbp respectively, so they have been dubbed “standard”, “ultra long” (UL), and “ultra ultra long” (UUL) for convenience. Nanopore datasets vary considerably in length characteristics, so cumulative length distributions are plotted for proper comparison (Figure 2b).

**Figure 2:**
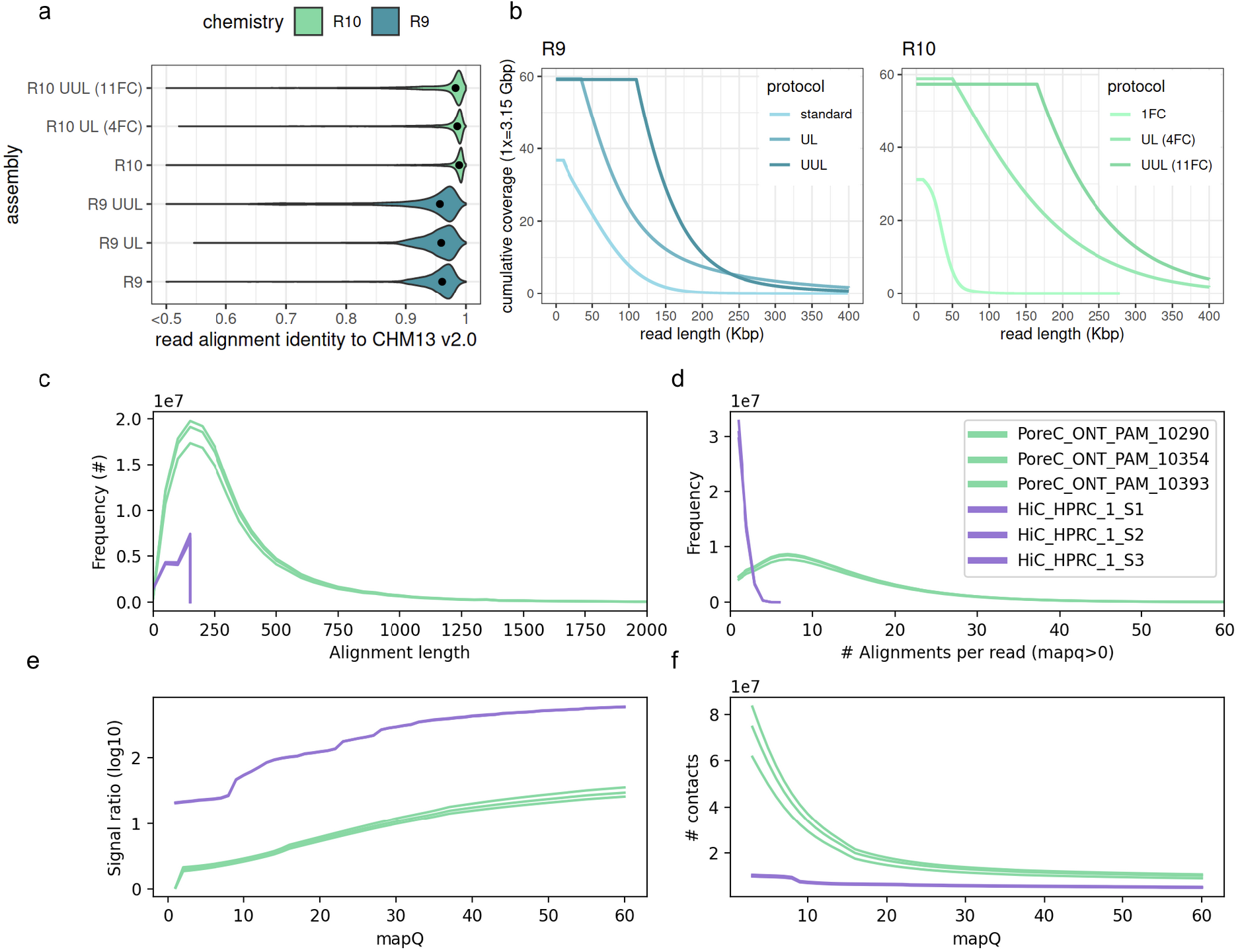
**a-b)** Identity and length metrics for nanopore read sets used in the HG002 evaluation. **c-f)** Pore-C and Hi-C metrics for contacts and signal ratio, measured on a per-library basis. “Alignment length” and “alignments per read” are proxies for subread statistics. Only mappings which are usable for phasing are shown, i.e with mapping quality (mapQ) >0 in a diploid reference. Signal ratio is computed using a high quality trio-phased assembly to indicate the number of consistent and inconsistent contacts (see methods).

For R10, a similar series of length distributions are used, with one key difference: the lowest coverage assembly used only a single PromethION flowcell (Figure 2b). For that dataset, reads were sheared to maximize throughput. In the R10 UL dataset, 4 flowcells of unsheared DNA were combined for a high coverage, longer dataset. Finally, in an attempt to maximize contiguity and find the limits of our methods with this data, we have also assembled a dataset containing only reads longer than 165Kbp, from 11 flowcells.

### Pore-C and Hi-C sequencing

An early version of the Pore-C protocol was performed and provided by ONT for use in this work, and compared to existing Hi-C datasets from the HPRC Year 1 data freeze. Notably, these Pore-C libraries do not have a larger modal alignment length than Hi-C (Figure 2c). The difference between Pore-C’s multi-contact concatemers and Hi-C’s 1-to-1 paired ligation are shown using alignments. Pore-C concatemers are composed of many smaller reads, or “subreads”, analogous to Hi-Cs paired reads. As a result of having many subreads, each Pore-C read has many alignments, and its contacts accumulate in an all-vs-all manner among subreads. The number of subreads (usable for phasing) can be in the dozens (Figure 2d). In contrast, Hi-C accumulates at most one long range contact per pair of reads, and in many cases, read pairs contain unmappable reads, which drastically reduces its throughput.

To demonstrate the practical differences between Pore-C and Hi-C that are particularly relevant to phasing, these results also show the signal ratio (Figure 2e) and the total number of contacts (Figure 2f) for reads which map to a diploid reference. Since mapping quality is used to filter contacts during phasing, the spectrum of signal ratio and number of contacts is plotted across observed map qualities. Signal ratio is computed using a trio-phased diploid reference, to estimate the number of consistent and inconsistent contacts (cis or trans w.r.t. haplotypes). In summary, contacts from Hi-C consistently have a higher signal ratio, while Pore-C produces more contacts.

### Assembers evaluated

For phasing and assembly quality evaluation, we compare to widely-used pipelines for diploid assembly that use ONT reads, PacBio CCS, or both. For PacBio CCS and hybrid CCS/ONT assemblers we have included Hifiasm and Verkko (Table 1). As the name implies, Hifiasm is an assembler for PacBio CCS (HiFi) data which has built-in methods for trio and Hi-C phasing. When comparing Shasta to Hifiasm the relative strengths of PacBio and nanopore become apparent. GFAse is also compared to Hifiasm’s native phasing methods to evaluate GFAse’s performance. Verkko is used in this comparison as an upper limit, since it uses both high coverage CCS and ONT reads to generate its assemblies. In this comparison, Hifiasm uses 30x coverage CCS and 30x coverage HiC. Verkko “production” assemblies use 42x CCS and 70x ONT >100Kbp, and trio Illumina. The “full coverage” Verkko assembly uses 190x CCS.

**Table 1:** Coverage summaries for HG002 assemblies evaluated in this analysis

|  |  | ONT |  | CCS |  | HiC / PoreC |
| --- | --- | --- | --- | --- | --- | --- |
| Assembler | Label | Coverage | N50 (Kbp) | Coverage | N50 (Kbp) | Coverage |
| HapDup (Shasta) | " | 38 | 31 |  |  |  |
| Shasta | R9 Standard | 37 | 60 |  |  | 45/30 |
| Shasta | R9 UL | 60 | 80 |  |  | 45/30 |
| Shasta | R9 UUL | 60 | 150 |  |  | 45/30 |
| Shasta | R10 (1FC) | 26 | 32 |  |  | 45/30 |
| Shasta | R10 UL (4FC) | 60 | 130 |  |  | 45/30 |
| Shasta | R10 UUL (11FC) | ~60 | 230 |  |  | 45/30 |
| Hifiasm Trio | " |  |  | 30 | 17.5 |  |
| Hifiasm Hi-C | " |  |  | 30 | 17.5 | 30 |
| Verkko Trio | production | 185.77 | 81.20 | 42.66 | 14.75 |  |
| Verkko Trio | full coverage | 971.71 | 50.65 | 169.03 | 17.22 |  |

For nanopore, HapDup has recently developed a combination of alignment-based and assembly-based methods for phasing long reads^22^. HapDup uses a linear unphased Shasta assembly as a starting point, phases aligned reads, and then generates a reference-free consensus for each haplotype. HapDup is specialized for structural variant detection in low coverage nanopore datasets, so it is a natural comparison point for phased Shasta assemblies.

### Phasing results

To evaluate phasing accuracy, the assemblies presented are aligned to a common reference and their heterozygous alleles are compared using WhatsHap to an orthogonally phased truth set, produced by NIST’s Genome In A Bottle^23^ consortium. Switch rate indicates how often alleles in the sample switch phase relative to the truth set, and hamming rate indicates the proportion of switched loci. Genotypes with allele sequences that do not both exactly match the reference VCF are not evaluated for phasing, which is accounted for by reporting the number of variants covered.

GFAse Trio uses k-mers from parental Illumina short reads to phase the heterozygous bubbles in the child Shasta assembly graph. Phasing results show that Shasta + GFAse Trio using Illumina reads outperforms Hifiasm Trio and Hifiasm HiC in terms of median switch rate (~ 0.0005) and hamming error (~ 0.0005). Shasta + GFAse trio results are also within range of the Verkko Trio ‘production’ assembly that uses both ONT and PacBio input reads. Two chromosomes, Chr15 and Chr16, have higher hamming rates across all the R10 Shasta assemblies, which may be due to the difficulty of resolving these chromosomes and identifying inversions relative to the hg38 reference. This method requires short read sequencing of both parents which, for various reasons, may not always be feasible or cost effective. For the more contiguous Shasta assemblies, trio results closely match the phasing results from Hi-C and PoreC phasing which don’t require any parental sequencing.

Shasta + GFAse assemblies consistently outperform native Hifiasm HiC phasing, and in some cases, hamming rates match or beat trio-phased Hifiasm. When comparing the total number of assessed variants, R10 assemblies drastically improve on R9, likely as a result of its lower error rate. This effect can be seen by comparing the standard length results to UL, in both R9 and R10. It is also evident from the HapDup results that polishing has a drastic effect on covered variants, since it uses a single flowcell R9 Shasta assembly. UL R10 assemblies reach nearly as many variants covered as Verkko while also producing comparable switch and hamming rates.

Yak Trio eval, an orthogonal evaluation method that uses parental short read k-mers to evaluate phasing accuracy, reports an order of magnitude higher switch and hamming error for Shasta assemblies compared to Verkko (Supplementary Table 2). This could be from a combination of greater genome coverage and systematic false positives in the switch analysis. As one possible explanation for the discrepancy between yak and WhatsHap, we observed that some of the yak switch blocks occur in regions of the Shasta assembly that differ from Verkko only in homozygous variants (Supplementary Figure 4a) and this is particularly evident in chrX of the male HG002 assembly (Supplementary Figure 4d). As an additional source of enrichment of errors, we observe that some yak switches occur in repetitive regions that contain homozygous homopolymer errors (Supplementary Figure 4c). When comparing Shasta Dipcall VCFs directly to Verkko Dipcall VCFs we observe a variant coverage of 2.48M at a hamming rate of 0.0003, but the total variants from Dipcall differ significantly in assembly specific Indels (Supplementary Table 3).

As evaluated by WhatsHap, GFAse consistently produces a median chromosomal hamming error of <0.001% for Shasta assemblies, which is reduced from an average chromosomal hamming of 40-50% for non-globally phased assemblies. In the single flowcell, standard length R10 experiment, R10 produces shorter haplotype bubbles than R9, which are then phased together at the chromosome scale with proximity ligation data (Figures 3 and 4d). In the high coverage UL R10 assembly, a large portion of the variants exist in continuous haplotypes directly from the Shasta assembler (Figure 4e). Generally, the graphs with high input N50 reach trio-level phasing accuracy when phased with HiC. The highly fragmented input graphs such as the Shasta “standard” R9 or the Hifiasm PacBio graphs converge to a less optimal phasing.

**Figure 3:**
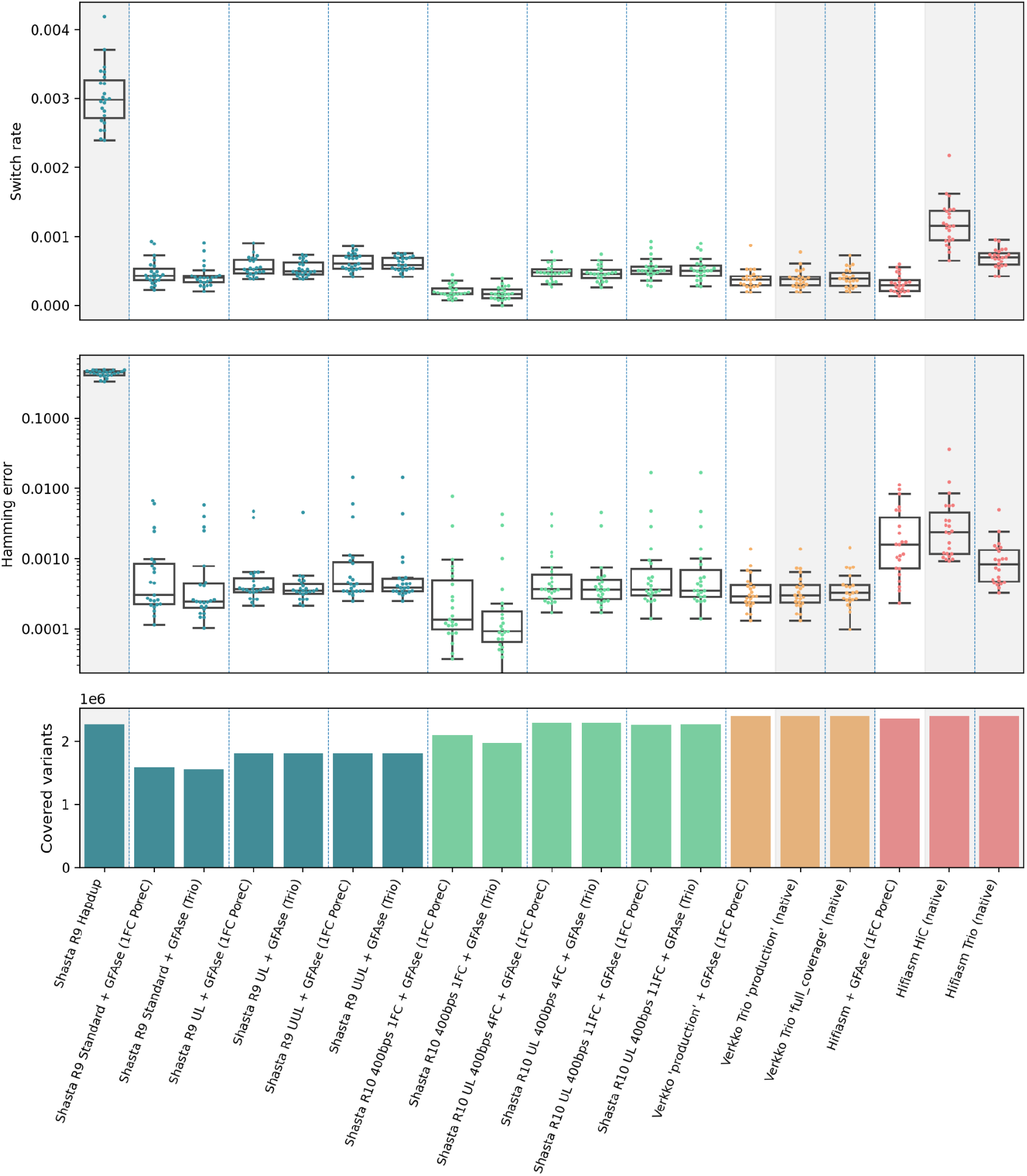
Phasing metrics for HG002 assemblies, as evaluated using the GIAB v4.2.1 benchmark VCF, phased with StrandSeq using WhatsHap (see methods). All shasta assemblies are unpolished. Assemblies not phased with GFAse are shaded gray. Each dot represents a chromosome error rate, generated by WhatsHap compare. Native Hifiasm HiC uses 30x coverage. Each pair of HiC is ~17x. PoreC flowcells have ~30x yield.

**Figure 4:**
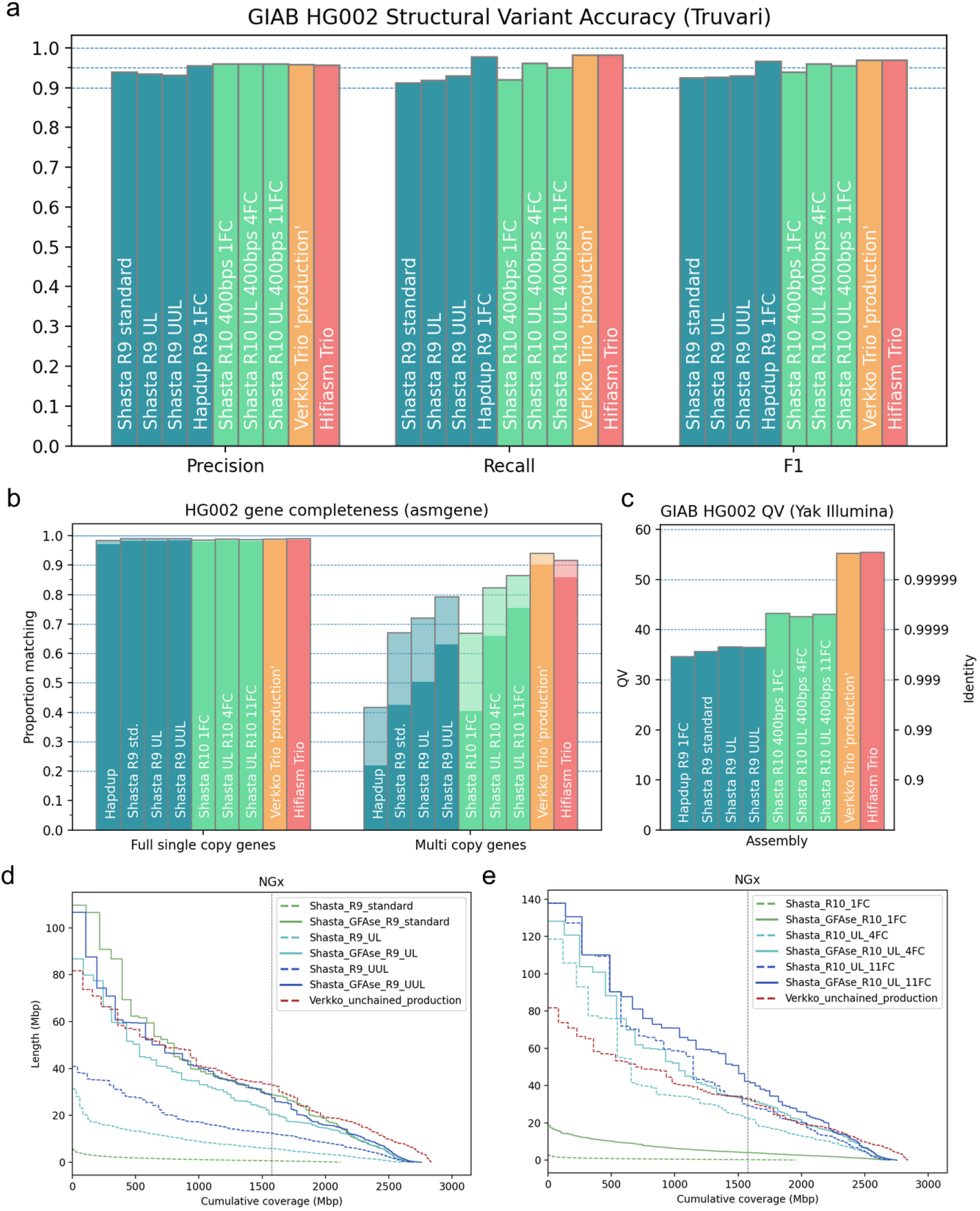
Structural variant, base level, and gene level accuracy metrics for HG002 assemblies. **a)** Base accuracy evaluated using yak with Illumina NovaSeq. **b)** Gene completeness measured by asmgene using human transcript sequences. “Full single copy” genes only indicate unfragmented, non-duplicated genes, matching transcripts by 99% or greater coverage and stratified by >97% (translucent) or >99% identity (opaque). Multicopy genes are similarly stratified. **c)** SVs evaluated using the GIAB Tier1 benchmark VCF with Truvari. **d-e)** NGx Plots for Shasta haplotypes, before and after unzipping bubble chains with GFAse. For comparison, the phased portion of the un-chained Verkko ‘production’ assembly is shown. The vertical line indicates the NG50 for each assembly.

**Figure 4:**
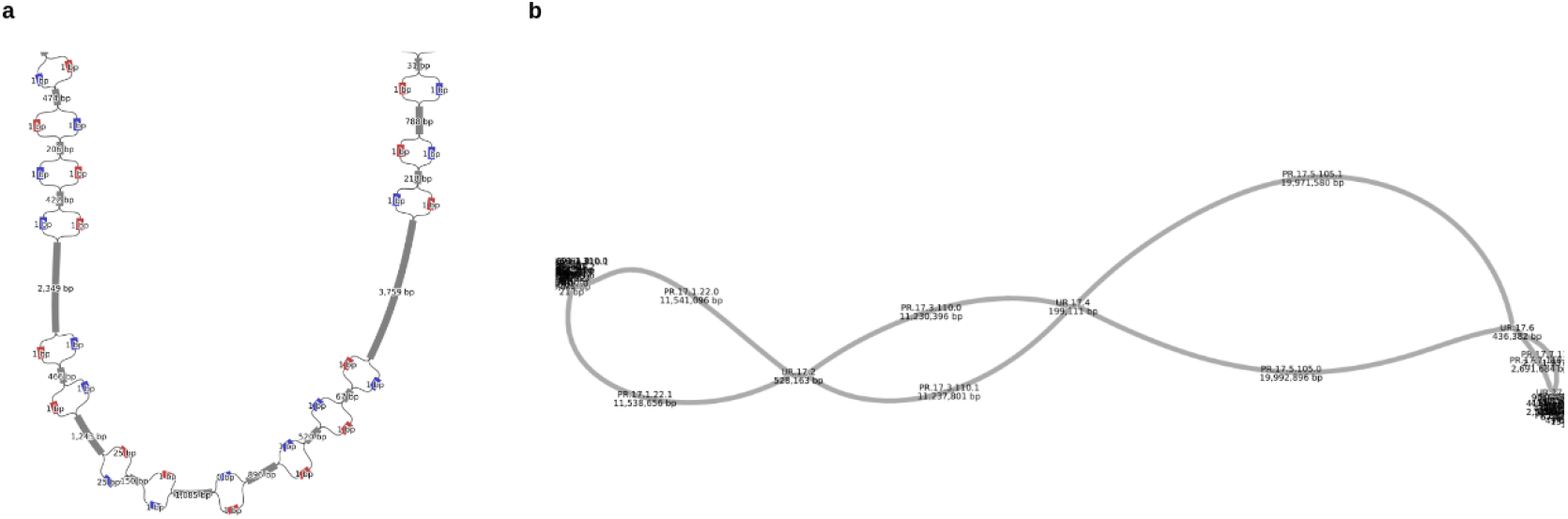
The two types of Shasta output graphs, visualized as a 2D layout in Bandage^31^ at two different scales. **a)** A subregion of the “Assembly-Detailed.gfa”, showing a near-variant scale nodes in a bubble chain and their phasing indicated by colors produced by Shasta. **b)** A subregion of the “Assembly-Phased.gfa” showing a phased portion of chr11 from HG002 which terminates at two tangles, presumably caused by telomeric and centromeric sequences.

In addition to this analysis, a limited number of HG005 R9 and R10 assemblies were evaluated (see Supplementary Figure 2 and Table 1), along with 4 HPRC R9 assemblies (Supplementary Figure 3). A similar trend in phasing accuracy is observed across all additional individuals, with the caveat that there is no StrandSeq-derived truth set for the HPRC assemblies.

By comparing Pore-C and Hi-C (Supplementary Figure 1) it is evident that more information is provided by one library of Pore-C than Hi-C. It takes 3 pairs (6×2 lanes, or ~45x) of Hi-C to reach a phasing that is roughly equivalent to 1FC Pore-C (30x). This is likely due to the higher number of total contacts per Pore-C read. In combination with the single-flowcell R10 assembly, we achieve an accurately phased human assembly with a total of 2 PromethION flowcells.

### Assembly quality

Structural variants were evaluated with Truvari and the GIAB HG002 Tier1 structural variant collection as a truth set. Shasta nanopore assemblies consistently yield structural variant F1 scores above 90%, with a peak score of >95% in the UL 4FC and 11FC R10 assemblies. Most notably, a single flowcell of R10 400bps nanopore data can reach an F1 score of around 94% (Figure 4a). HapDup^24^, which starts with Shasta assembly, uses a local realignment and polishing to recover short collapses, which is a probable cause for its greater recall. Precisions from R10 assemblies match or exceed HiFi-based methods, but recall values do not, most likely as a result of collapse in the repetitive regions of the genome.

A similar trend is seen in the gene level analysis performed by asmgene (Figure 4b), in which the number of full single copy genes are comparable to HiFi or hybrid assemblies, but multi copy genes are reduced in comparison. For gene completeness, the highest scoring Shasta assembly reached ~75% multicopy genes at a 99% identity threshold, which is now 10% lower than Hifiasm. In terms of base level quality, R9 assemblies have base qualities greater than Q30, while R10 assemblies are greater than Q40 and PacBio HiFi or hybrid assemblies exceed Q50 (Figure 4c).

Shasta assemblies are chained and unzipped by GFAse to achieve greater continuity. In theory, only 0.1% of the assembly would need to be assembled as diploid bubbles to produce a fully phased assembly after unzipping, but in practice, mapping and optimization with proximity ligation reads is easier when haplotypes are long. Before chaining, bubble N50s range from <1Mbp in the low coverage experiments up to 39.7Mbp in the highest quality assembly. For the various R9 assemblies, the chained and unzipped assemblies remain consistent in length despite their variable input lengths, which is shown by their largely overlapping post-GFAse NGx distributions (Figure 4d–4e). R10 sequencing protocols are currently limited by throughput, so sheared reads were used for the single flowcell experiment, which do not enable the same level of contiguity as the UL assemblies. However, for the higher cost 4FC and 11FC UL experiments, the upper limit on contiguity exceeds our previous most contiguous R9 assemblies.

### Non-human assemblies

To test Shasta’s phased assembly methods outside the context of human genomes and basecalling, two species were assembled: the dwarf cuttlefish (*Sepia bandensis*) and the broad bordered yellow underwing moth (*Noctua fimbriata*). *S. bandensis*, previously unsequenced by long reads, was sequenced for this work to a depth of ~105x with 30x >100Kbp length. *N. fimbriata* was previously sequenced by the Darwin Tree of Life^25^ to a depth of 87x with an N50 of 28.7Kbp.

In non-human assemblies, truth sets are limited, so this analysis relies on BUSCO to evaluate gene completeness and extent of phasing (Table 2). By comparing the BUSCO score for the full diploid assembly, and one haplotype of the diploid assembly, the number of phased and unphased genes is measurable. In both species presented, complete phased genes are estimated to range from 86% to 89% of the 954 genes in the metazoan dataset.

**Table 2:** Non-human Shasta phasing metrics, in terms of BUSCO gene completeness. These results use the metazoan dataset, which has 954 genes.

|  | Complete and single-copy (S) |  | Complete and duplicated (D) |  |  |  |  |
| --- | --- | --- | --- | --- | --- | --- | --- |
|  | Both haps | One hap | Both haps | One hap | Total diploid | Total Haploid | Diploid N50 |
| ***Sepia bandensis*** | 5.97% | 95.39% | 90.15% | 0.73% | 5330371146 | 415993994 | 3794550 |
| ***Noctua fimbriata*** | 7.02% | 93.29% | 91.40% | 4.82% | 488722705 | 21756252 | 5094850 |

### Resource usage

For the slowest assembly evaluated (4 flowcell R10 UL), Shasta runs in 12 hours on a 64 thread 1.2TB AWS instance. This allows for 14 assemblies to be run in the same amount of core hours as a single Verkko assembly (Table 3). For unfragmented assemblies, GFAse has variable run time of 2.3hr to 4.6hr using 64 threads, which depends on the number of contacts in the alignments. In the worst case scenario, with a highly fragmented GFA such as the unphased Hifiasm graph, and high contact dataset such as PoreC, GFAse can take up to 12 hours.

**Table 3:**
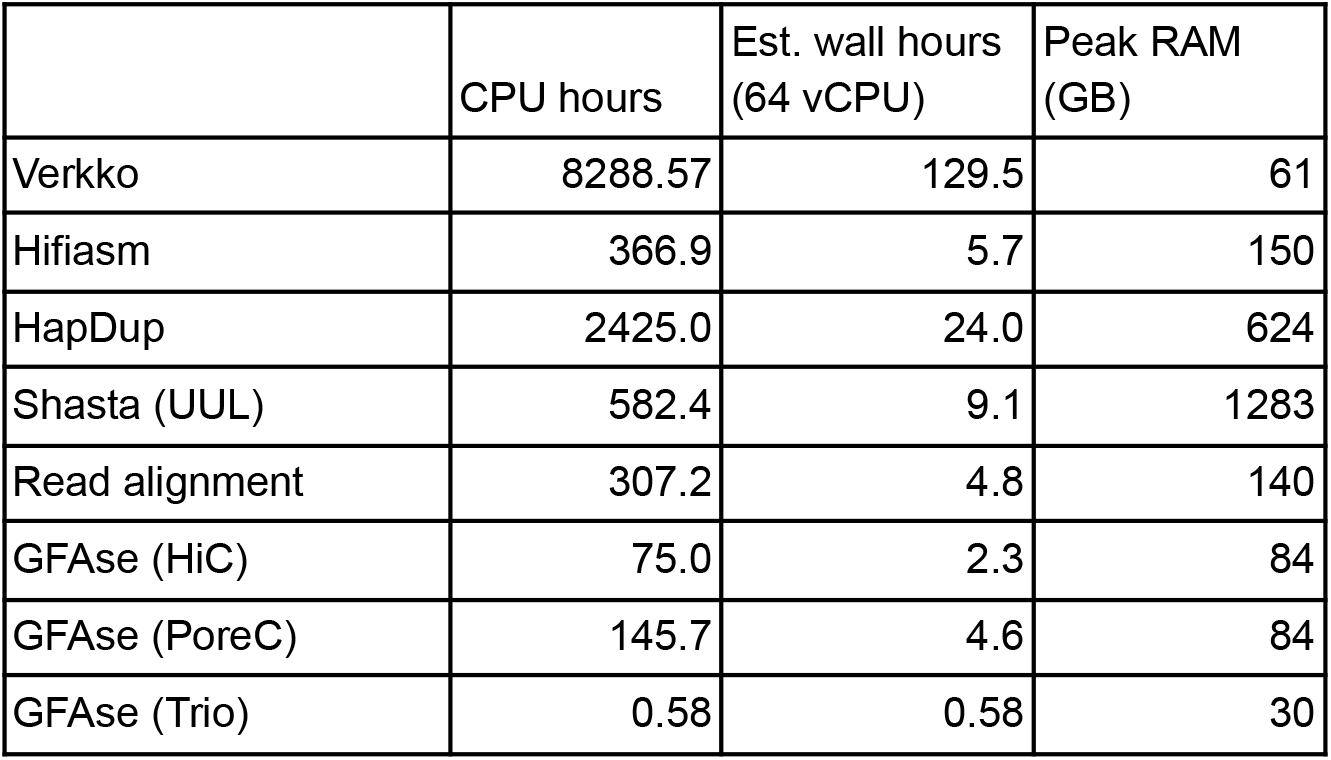
Run time performance for various assemblers presented in this paper. Separate times shown for GFAse HiC and PoreC as a result of its dependence on the number of contacts. For HapDup, CPU hours and wall hours show the sum of multiple steps in the HapDup pipeline, including an initial Shasta assembly, which used 96 cores. GFAse trio runs single threaded.

|  | CPU hours | Est. wall hours<br>(64 vCPU) | Peak RAM<br>(GB) |
| --- | --- | --- | --- |
| Verkko | 8288.57 | 129.5 | 61 |
| Hifiasm | 366.9 | 5.7 | 150 |
| HapDup | 2425.0 | 24.0 | 624 |
| Shasta (UUL) | 582.4 | 9.1 | 1283 |
| Read alignment | 307.2 | 4.8 | 140 |
| GFase (HiC) | 75.0 | 2.3 | 84 |
| GFase (PoreC) | 145.7 | 4.6 | 84 |
| GFase (Trio) | 0.58 | 0.58 | 30 |

## Discussion

In this work, we use the latest advances in nanopore sequencing to simplify the challenge of producing phased *de novo* assemblies. We demonstrate accurate phasing in a two PromethION flowcell pipeline, using one long read flowcell and one Pore-C flowcell. Phased contiguities yielded by our higher coverage experiments are unmatched in previous nanopore assemblies, and Pore-C as a phasing data type is unexplored in prior publications. The tools presented are efficient and modular, and they rely on sequencing protocols which have a relatively short turn around^26^, enabling rapid prototyping.

We focus on nanopore sequencing because, despite its currently lower base-level accuracy relative to competitors, its ability to sequence native DNA means that its upper read length is essentially limited only by the library preparation and loading procedure, which gives it a unique advantage over methods which rely on synthesis or amplification^27^. With recent changes in chemistry and computational methods, this tradeoff in accuracy has reduced, while cost and throughput have improved. In three years, since Shafin et al 2020, R9 median accuracy has increased from 90% to 95%, effectively reducing error by half. In the same time span, protocols for nanopore library preparation have improved average N50s from 42Kbp to >100Kbp, more than doubling. Now, with the R10.4.1 chemistry in production, we see yet another reduction in error, bringing accuracy into the 98-99% range.

The alternative sequence type to nanopore, PacBio HiFi (CCS), has accuracies of >99.9%^28^, but its reads are size-selected, ranging from 10Kbp to 30Kbp. This means that they have the accuracy to distinguish copies of less diverged repetitive units in the genome, but not necessarily the length to span them. Hybrid assemblers have integrated both ONT and HiFi data to accommodate for this, in addition to parental (“trio”) Illumina data or Illumina proximity ligation data (Hi-C) to assist with phasing. Hybrid methods have achieved unprecedented accuracy and contiguity by leveraging the strengths of 3-4 different sequencers, but as a result they are also costly in terms of resources, logistics, and time.

Using technological advances in ONT sequencing, our results produce notable improvement over the previous standards for nanopore phasing. Evaluation metrics for phased assembly are approaching that of the hybrid assemblers in gene completeness, contiguity, and phasing accuracy, showing promise for future single sequencer pipelines. When considering that all Shasta assemblies generated for this paper are unpolished, it is clear that there is room for further improvement. Contiguity and completeness are likely to continue to grow proportionally with additional throughput of long reads. To make full use of the longest reads, further method development is underway to address repetitive regions which defy the diploid assumption of our current methods. The consistent trajectory of ONT quality and throughput has motivated our work, and we aim to continue to adapt long read assembly methods to future improvements.

From a software development perspective, the tools presented in this paper are written with the goal of modularity, interpretability, and flexibility of usage. The fully specified graph outputs of Shasta make it an ideal resource for downstream development and analysis, as is demonstrated by the application of GFAse in this context. GFAse employs transparent and reusable data structures, and similar to Shasta, produces comprehensive outputs that describe the homology, proximity linkage, and inferred haplotype chains in the graph. Relying directly on alignments, GFAse is capable of using any data type for phasing which can be aligned to the assembly in BAM format. In theory, long reads, or even conventional linked reads could be used as phasing information if their alignments span the unphased regions of an assembly. This makes it a flexible module for future applications, as long range linked data types continue to evolve.

## Methods

### Shasta - *De novo* phasing with nanopore reads

Shasta is an assembler specialized for nanopore reads, and it uses the Overlap Layout Consensus paradigm of assembly. It starts by reducing reads into vectors of fixed length subsequences or markers, and then it computes an approximate overlap among reads using a variation of the MinHash algorithm^29^. Shasta refines its candidate overlaps using alignment in marker space, and reduces the overlap graph by filtering alignments and creating a k-nearest neighbor graph. For more details, see the initial 2020 publication, or the online documentation^26,30^.

In this updated version, Shasta performs *de novo* phasing internally, using only conventional nanopore reads. Shasta uses a data structure referred to as a “phasing graph”, built from a marker representation of the reads. The phasing graph is created following the overlap stage of assembly, and it describes the coverage in terms of read IDs covering each branch of a heterozygous diploid bubble found in a graph. For each pair of bubbles, a Bayesian model computes the probability that they are either uncorrelated or in one of two possible phased orientations with respect to each other. By iteratively aggregating bubbles with this Bayesian criteria for correlation, groups of phased bubbles are established, while uncorrelated error bubbles remain isolated.

Given a set of phased bubbles or a “component”, Shasta then identifies local bubble chains within each set, which are bubbles in series, constituting collinearly traversable regions of the graph^30^. These bubble chains have a strict topology in which elements in the chain can either be homozygous unphased nodes or heterozygous phased pairs of nodes (see Figure 4A). Bubble chains are the basis for subsequent unzipping into haplotypes. For completeness, Shasta generates output GFAs containing the succinct representation of edits, or “Detailed” graph, as well as the larger unzipped haplotype representation, or “Phased” graph. Both of these representations contain bubble chains with differing length haplotype sequences. The “Phased” representation is convenient for downstream phasing because its sequences tend to greatly exceed the length of a read and multiple variants can be spanned with conventional mapping. On the other hand, the “Detailed” representation summarizes the edits between phased haplotypes, and represents longer scale phasing using a path in the GFA. Graphs generated by Shasta contain “blunt” or non-overlapping nodes, which makes chaining them trivial.

### GFAse - Phasing graphs with proximity ligation data

GFAse relies on conventional mappings for phasing information. HiC, PoreC, or other proximity-ligated reads are mapped to the GFA contigs using whichever mapper is most appropriate for the sequence type. The strength of the proximity linkage between any two contigs is updated as a sum, in the form of a weighted edge in a “contact graph”, where the weight is the number of reads linking them. Uninformative mappings which do not cover a heterozygous site are filtered by setting a map quality threshold.

To phase the graph, GFAse first identifies haplotypic bubbles. Two methods are available in GFAse: assembler annotation and sequence similarity search. Efficient similarity search is accomplished with a variation of MinHash^29^ similar to that used by Shasta^26^, and then refined with full scale alignment with minimap2^32^.

Phases are optimized using a stochastic method which approximates a solution to the optimization variant of the max-cut problem^10,13,17^. The method depends on an objective function which penalizes inconsistent contacts and rewards consistent contacts. If any two bubbles are compared, there are four possible contacts, and only contacts linking the contigs in matching phases have positive scores. For GFAse, a variation on existing methods was introduced to improve reproducibility of the stochastic method and perform better on fragmented graphs in which the state space is much larger. In short, the method takes samples from repeated greedy optimizations of randomly initialized phase states and accumulates a distribution of orientations, which is then used to merge bubbles that are most consistent. One benefit of sampling many times with few iterations is that samples are independent and can be multithreaded.

**Figure 5:**
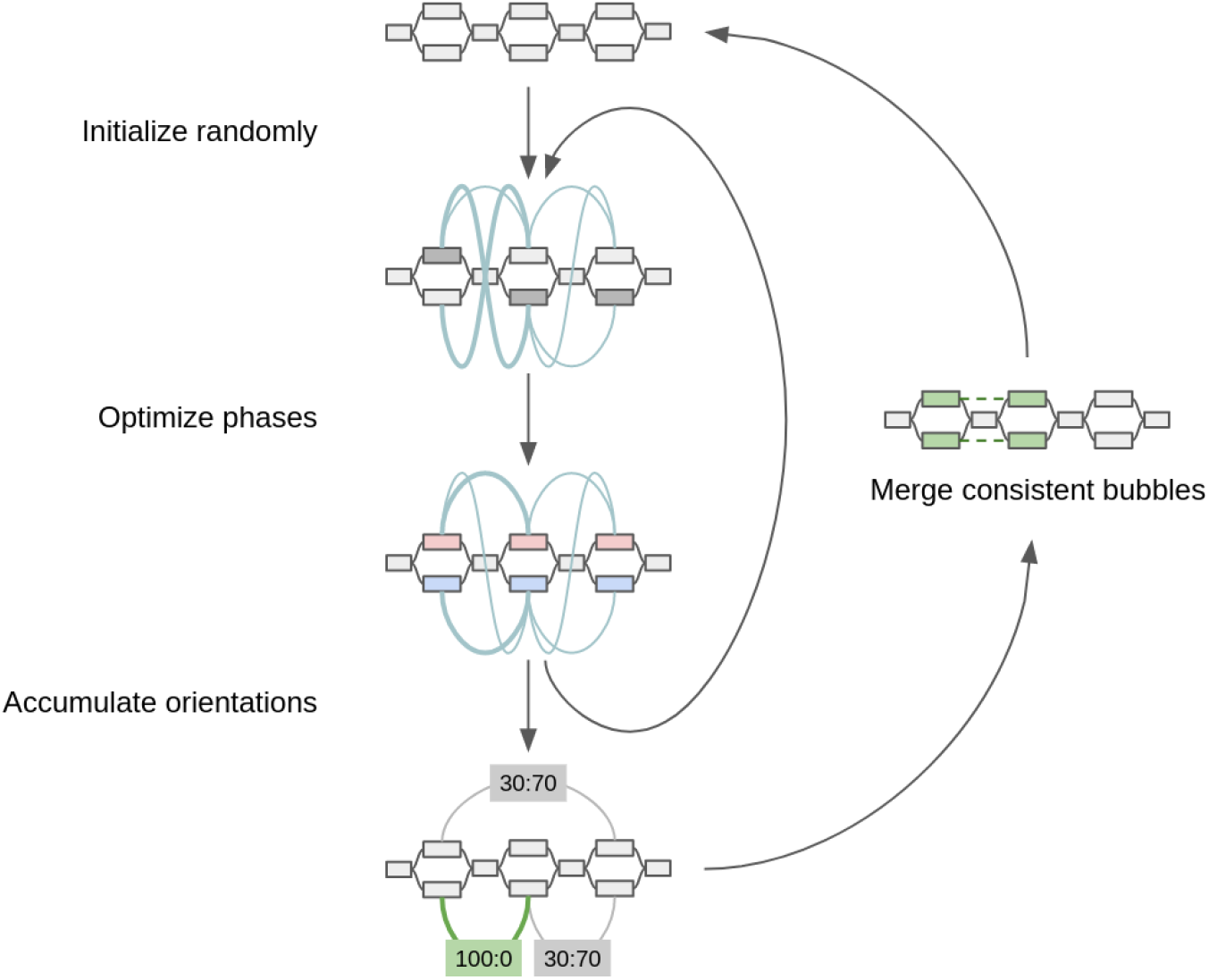
Diagram of sampling method for optimizing proximity linkages in an assembly graph. Edge weights in the contact graph are represented by teal curves. For each inner iteration, a greedily converged phase state is used to update a distribution of orientations among bubbles. Bubbles with the strongest signal at the end of sampling are merged for successive iterations. By the end of each round *r* of merging, the largest possible bubble set is 2^*r* in size.

### GFAse - Phasing with parental data

Homozygous parental k-mers are selected from each parent and used to phase the “detailed” assembly GFA by counting parental k-mers in the heterozygous bubbles. To process the parental sequence data, reads are broken into 31 base pair k-mers using kmc3^33^. Kmc3 subtract was used to identify k-mers that are unique to each parent. Finally, unique homozygous k-mers are matched to child k-mers on heterozygous bubbles assigning a phase to bubble components, using a simple majority vote. Illumina reads for the HG002 Ashkenazi Jewish Trio sample were obtained from the publically available 1000 Genomes Project (fc-4310e737-a388-4a10-8c9e-babe06aaf0cf/working/HPRC_PLUS/HG002/raw_data/Illumina/p arents/HG003 and fc-4310e737-a388-4a10-8c9e-babe06aaf0cf/working/HPRC_PLUS/HG002/raw_data/Illumina/pa rents/HG004).

### GFAse - Chaining phased graphs

With a phased assembly graph, adjacent bubbles are chained in a manner similar to scaffolding, to extend haplotypes. GFAse first loads the GFA using the VG HandleGraph data structure^34^ and identifies tractable regions as anything which follows a strict diploid bubble chain topology. Diploid nodes have exactly one two-hop neighbor and, at most, two direct adjacencies in each direction. Chains are then identified by traversing contiguous subgraphs of labeled nodes. With bubble chains identified, haplotypes are labeled with paths in the GFA formalism. Then they can be “unzipped” trivially by traversing them and duplicating the homozygous nodes into both haplotypes. The strict definition of bubble chains used in this method is intended to maximize fidelity to the input graph by reducing misjoins in the chaining step.

### Sequencing and data acquisition

#### R9 Standard

Sequencing was performed as described in Shafin et al. 2020, and re-basecalled with Guppy v5.0.7.

#### R9 UL

ONT data from the GIAB consortium^35^ was re-basecalled with Guppy >v5.0 and combined with the R9 Standard dataset to provide longer reads.

#### R9 UUL

DNA extractions from 6 million cells of HG002 were prepared using Circulomics Nanobind CBB Kit (Pacific Biosciences, 102-301-900). Libraries were prepared using Ultra-Long DNA Sequencing Kit (SQKULK001). The libraries were sequenced on Flowcell R9.4.1 on PromethION for 72 hours. Flowcells were washed using the Flowcell Wash Kit (EXP-WSH004) every 24 hours. Fresh libraries were loaded after each wash.

#### R10

DNA extractions from 5 million cells of HG002 were prepared using PacBio Nanobind CBB kit (SKU 102-301-900) according to UHMW DNA Extraction Cultured Cells protocol (EXT-CLU-001). Standard pipette tips were used to generate a homogenous sample before DNA shearing. DNA was sheared to a target size of 50kb on Megaruptor^®^ 3. Samples were normalized to 100 μL volume at 50 ng/μL concentration and sheared at speed 27 using Megaruptor shearing kit (E07010003). DNA size was assessed post shearing on Agilent Femto Pulse System using gDNA 165kb Analysis Kit (FP-1002-0275). Post shearing, DNA size selection was performed using PacBio SRE kit (SKU 102-208-300) following manufacturer’s recommendations. Libraries were prepared using Oxford Nanopore Ligation Sequencing Kit V14 (SQK-LSK114) according to Oxford Nanopore protocol GDE_9161_v114_revK_29Jun2022. The libraries were sequenced on Flowcell R10.4.1 on PromethION for 96 hours. Flowcells were washed using the Flowcell Wash Kit (EXP-WSH004) every 24 hours. Fresh libraries were loaded after each wash.

#### R10 UL

Nanopore sequencing dataset were generated following Oxford Nanopore protocol ULK_9177_v114_revC_27Nov2022. DNA extractions from 6 million cells of HG002 were prepared using Monarch^®^ HMW DNA Extraction Kit for Tissue (New England Biolabs, T3060). Libraries were prepared using Ultra-Long DNA Sequencing Kit (SQKULK114). The libraries were sequenced on Flowcell R10.4.1 on PromethION for 72 hours. Flowcells were washed using the Flowcell Wash Kit (EXP-WSH004) every 24 hours. Fresh libraries were loaded after each wash.

##### Sepia bandensis

Testes tissue (3-5 mg) from an adult male dwarf cuttlefish was homogenized in PBS using a Dounce homogenizer. This was followed by DNA extraction using Circulomics Nanobind Tissue Kit (Pacific Biosciences, 102-302-100). Libraries were prepared using Ultra-Long DNA Sequencing Kit (SQKULK001). The libraries were sequenced on Flowcell R9.4.1 on PromethION for 72 hours. Flowcells were washed using the Flowcell Wash Kit (EXP-WSH004) every 24 hours. Fresh libraries were loaded after each wash.

##### Noctua fimbriata

ONT data was acquired from the Darwin Tree of Life project^25^.

### Generating assemblies

To generate nanopore assemblies, Shasta (v0.10.0 unless otherwise specified) was run with the appropriate configuration for each data type, as follows:

#### R9 Standard

--config Nanopore-Phased-May2022
--Reads.minReadLength 10000
--Assembly.mode2.phasing.minLogP 30

#### R9 UL

--config Nanopore-UL-Phased-May2022
--Reads.minReadLength 10000
--Reads.desiredCoverage 180000000000
--Assembly.mode2.phasing.minLogP 50

#### R9 UUL

--config Nanopore-UL-Phased-May2022
--Reads.minReadLength 110000
--Assembly.mode2.phasing.minLogP 50

#### R10 (1FC)

--config Nanopore-Phased-R10-Fast-Nov2022
--Assembly.mode2.phasing.minLogP 20

#### R10 UL (4FC)

Configs for *Nanopore-Phased-R10-Fast-Nov2022* and *Nanopore-UL-Phased-May2022* were merged, with *Nanopore-Phased-R10-Fast-Nov2022* taking precedence for any conflicting parameters. The following parameters were then added:

--Assembly.mode2.phasing.minLogP 20
--Reads.minReadLength 50000

#### R10 UUL (11FC)

##### (Shasta v0.11.1)

--config Nanopore-Phased-R10-Fast-Nov2022
--Kmers.probability 0.05
--MinHash.minBucketSize 20
--MinHash.maxBucketSize 60
--Align.minAlignedMarkerCount 2500
--Reads.minReadLength 170000

###### Sepia bandensis

--config Nanopore-UL-Phased-May2022
--Reads.desiredCoverage 400G

###### Noctua fimbriata

--config Nanopore-Phased-May2022

### Evaluation

Input data was evaluated using an alignment based QC tool called WamBam, which iterates BAMs and produces read identity and read length stats^36^. PoreC statistics were generated via a similar method using the “evaluate_contacts” executable provided in the GFAse repository. Reads were aligned to both haplotypes of the trio phased Verkko “full coverage” HG002 assembly, and statistics were calculated by accumulating loci and lengths for each mapping of each read ID. For the “signal ratio” calculation, a contact map was constructed by building a graph similar to that used in phasing methods. Any two mappings of the same read ID constitute an edge, and they are binned by the minimum mapping quality of the pair. Edges that cross from one haplotype to another are considered inconsistent with the true phasing, and are used to compute a ratio of consistent to inconsistent edges.

As a truth set for phasing, chromosome-length haplotypes were generated using Strand-seq and long reads. In order to generate haplotypes, we have used a combination of Strand-seq data and PacBio Hifi reads from the same individual (HG005 and HG002). Sparse and chromosome-length haplotypes were generated using Strand-seq data and R package StrandPhaseR (version 0.99) as previously described^37^. Next, we detected inverted regions using Strand-seq data and manually curated this list of inversions as previously described^38^. We have used this set of inversions to correct Strand-seq phasing over these regions with the StrandPhaseR function called ‘correctInvertedRegionPhasing’ as previously described^38^. After inversion phase correction, we generated dense chromosome-length haplotypes using a combination of Strand-seq haplotypes and PacBio long reads as previously described in the integrative phasing framework^37,39^. Integrative phasing was completed using WhatsHap (version 1.0)^9^. For integrative phasing, we used a defined set of variant positions (available at, https://ftp-trace.ncbi.nlm.nih.gov/ReferenceSamples/giab/release/ChineseTrio/HG005_NA24631_son/NISTv4.2.1/GRCh38/HG005_GRCh38_1_22_v4.2.1_benchmark.vcf.gz and https://ftp-trace.ncbi.nlm.nih.gov/ReferenceSamples/giab/release/AshkenazimTrio/HG002_NA24385_son/NISTv4.2.1/GRCh38/HG002_GRCh38_1_22_v4.2.1_benchmark.vcf.gz). To include indels into the final callset, we have run WhatsHap with --indels parameter.

To analyze phasing accuracy, a combination of Dipcall^40^ and WhatsHap was used. Dipcall is a reference-based variant caller. For a set of phased assemblies, it produces a VCF file of single nucleotide variants (SNVs) as well as small insertions and deletions (INDELs). For the male HG002 sample, Dipcall was run using the GRCh38 reference, but set to treat the PAR region as autosomal regions. Phase set tags were manually added to the Dipcall VCF file before being used by WhatsHap. WhatsHap “compare” assesses switch error and hamming distance in the phased assemblies by comparing the phasing of alleles in the NIST’s Genome In a Bottle^35^ truth set to the Dipcall VCF file. WhatsHap “compare” only includes variants in the analysis that have identical alleles in the truth and query VCF file, making it robust to SNVs caused by sequencing errors. WhatsHap “stats” was run with a “chromosme_lengths” input file to calculate phasing statistics.

Collapses and misassemblies within genes were evaluated using minimap2^32^, asmgene and the publically available Ensemble genes as input (https://ftp.ensembl.org/pub/release-87/fasta/homo_sapiens/cdna/Homo_sapiens.GRCh38.cdna.all.fa.gz). Ensemble cDNA was aligned to the CHM13 v2.0 reference and the HG002 assemblies with minimap2 using “-cx splice:hq” for intra-species cDNA alignment. Asmgene, a part of minimap2, selects the longest isoform from overlapping alignments and counts it as matching the reference if it covers >99% of the transcript length with a mapping identity above an input threshold. To account for ONT’s sequencing error profiles, asmgene was run with a mapping identity threshold of 97% instead of 99% used for HiFi assemblies. Transcripts are counted as single-copy if they uniquely align to the reference and multi-copy if they align to multiple loci.

Base quality was estimated using yak^41^, based on the k-mer content of Illumina short reads. Each phased assembly was evaluated separately. K-mer in the short reads were counted using “yak count -b 37”, and quality values (QV) were estimated using “yak qv -K 3.2g -l 100k”. For HG002, we used the 30x Illumina Novaseq PCR-free read set publically available at the Google bucket gs://deepvariant/benchmarking/fastq/wgs_pcr_free/30x/. For the 4 samples from the HPRC (HG01993, HG02132, HG02647, and HG03669), we used 30x Illumina short-reads from the high coverage readset of the 1000 Genomes Project samples^42^.

Non-human assemblies (*Noctua fimbriata* and *Sepia bandensis*) were evaluated using default arguments for BUSCO v5.4.3, and the “metazoan_odb10” dataset. To attempt to evaluate the number of phased genes, BUSCO was run twice: once with both haplotypes, and once with one haplotype. For the “both haplotypes” evaluation, the entire diploid assembly was provided to BUSCO. For the “one haplotype” evaluation, one of each haplotype from the phased regions was removed, and the remaining sequences were evaluated by BUSCO.

Switch error in the phased assemblies was also estimated from Illumina short reads from parents. We used yak to count the k-mer in the short reads, as above, and “yak trioeval” to compute the estimated switch error rate. As above, the read sets for HG002’s parents were downloaded from the same Google Bucket as for the base quality evaluation, and from the 1000 Genomes project’s dataset for the 4 HPRC samples.

Structural variants (SVs) were called from the phased assemblies using dipdiff, a modified version of the SVIM-ASM tool^43^. In HG002 assemblies, the SVs were called against GRCh37 and evaluated with the GIAB SV truthset^23^ using Truvari^44^. Truvari’s “bench” command was run with “--no-ref a -r 2000 -C 2000” to ignore missing homozygous calls for the reference allele, and match variants up to 2,000 bp away from each other.

## Acknowledgements

This work was funded in part by the National Institutes of Health under award numbers: R01HG010485, U24HG010262, U24HG011853, OT3HL142481, U01HG010961, and OT2OD033761. A portion of the R10 HG002 dataset labeled “UUL” in this work was supported by startup funds (Miten Jain, Genome Technology Laboratory, Northeastern University).

Cuttlefish Nanopore work was supported by Oxford Nanopore Technologies grant SC20130149 (awarded to Mark Akeson, UCSC Nanopore Group). We acknowledge the support of Oxford Nanopore Technologies staff in generating this data set, in particular the pore-C data.

