## Supplementary Figures and Tables for "Phased nanopore assembly with Shasta and modular graph phasing with GFAse"

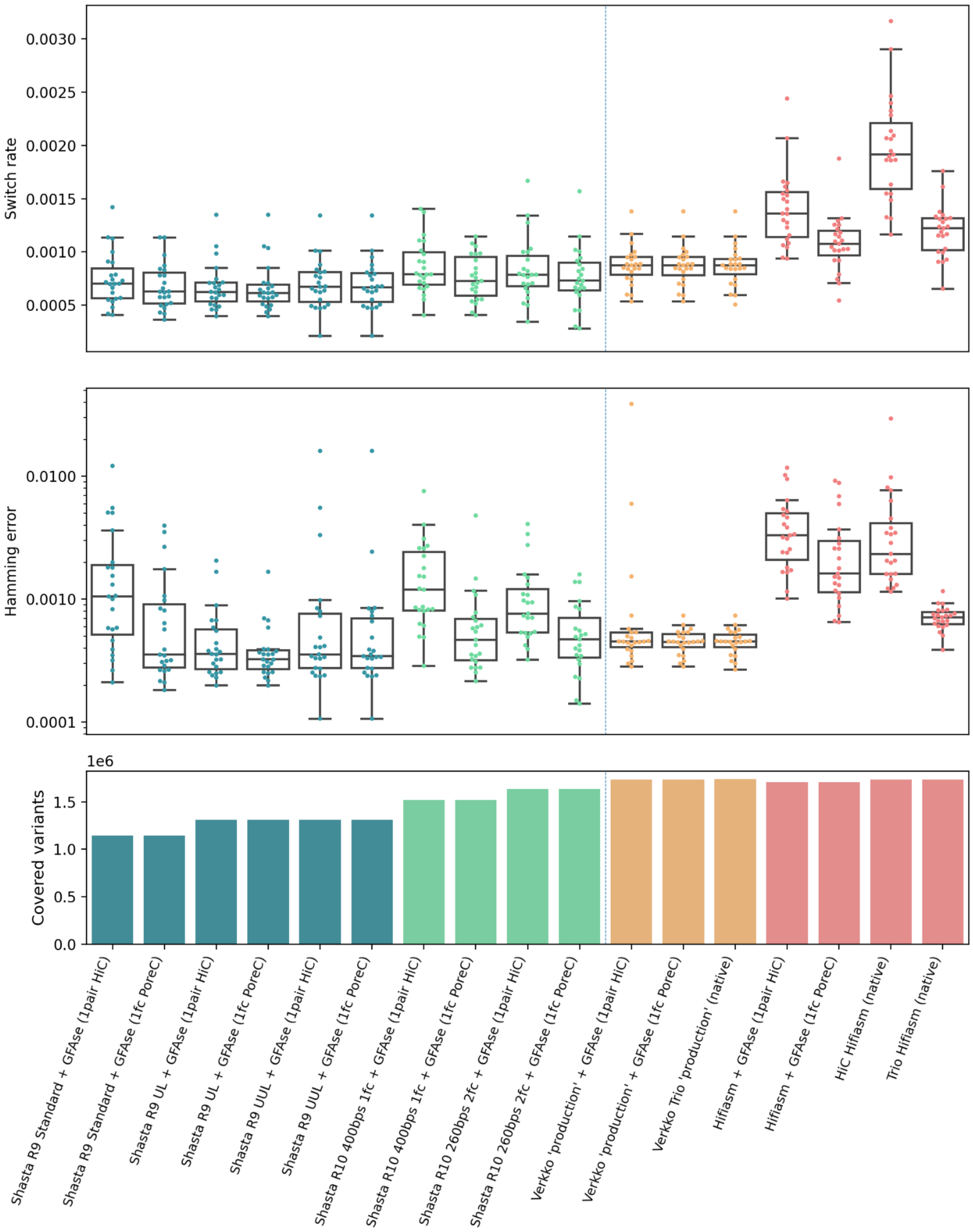


**Supplementary Figure 1:** Comparison of single pair HiC vs single flowcell PoreC (using GIAB StrandSeqANDTrio v4.2.1). Note that this truth set has fewer variants than that used in the main text.

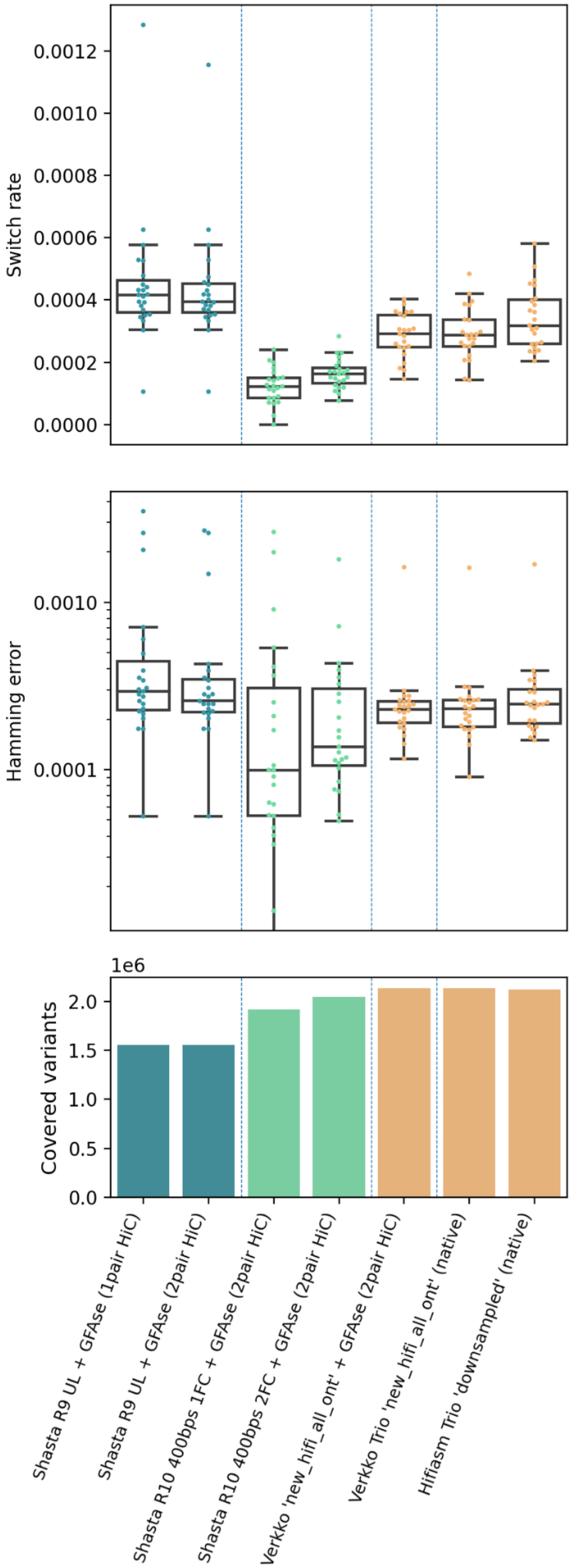


**Supplementary Figure 2:** HG005 evaluation using StrandSeq integrative phasing as a truth set.

| | | ONT | | CCS | | HiC |
|---|---|---|---|---|---|---|
| **Assembler** | **Label** | **Coverage** | **N50 (Kbp)** | **Coverage** | **N50 (Kbp)** | **Coverage** |
| **Shasta** | R9 UL | 57.1 | 111 | | | 30 |
| **Shasta** | R10 1FC | 34.0 | 32 | | | 30 |
| **Shasta** | R10 2FC | 57.2 | 34 | | | 30 |
| **Verkko Trio** | new_hifi_all_ont | 185.8 | 81.2 | 33.7 | 16.06 | 30 |

**Supplementary Table 1:** HG005 assembly input data

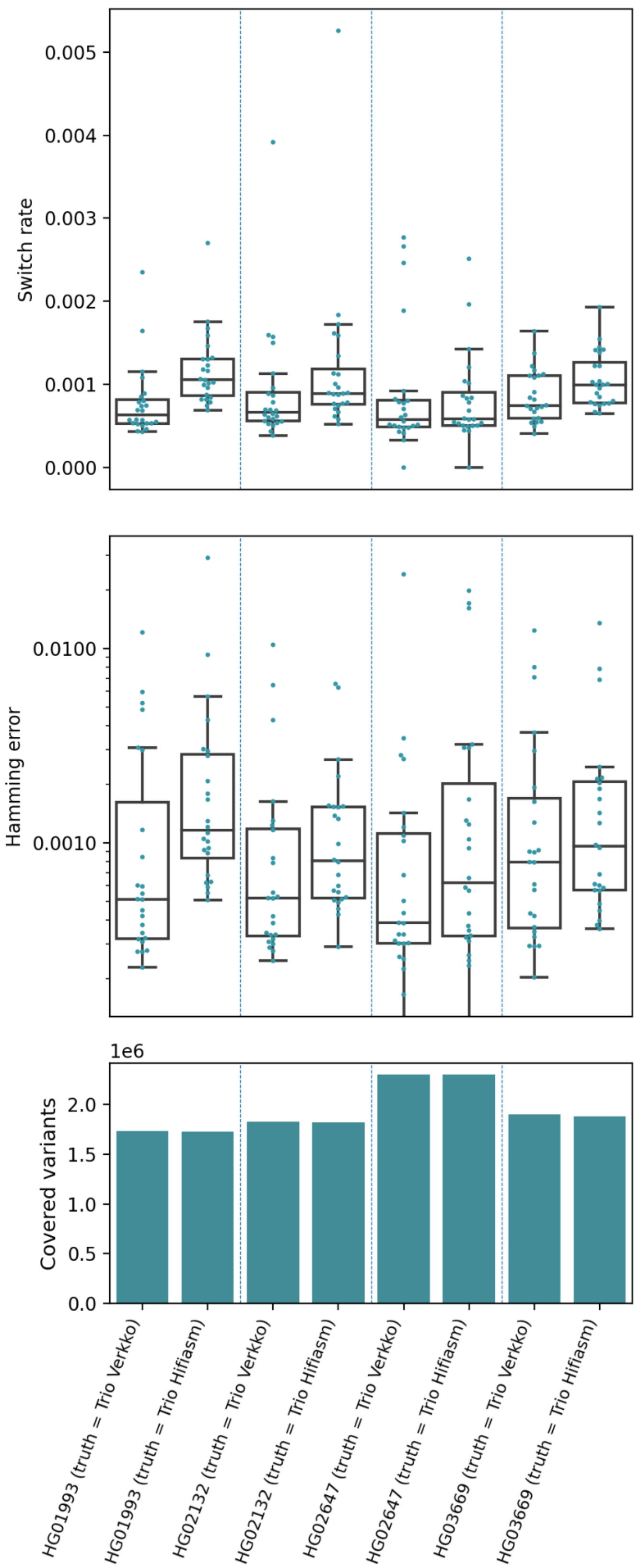


**Supplementary Figure 3:** HPRC evaluation using Trio Hifiasm and Trio Verkko aligned to HG38 as truth.

| assembly | switch_rate | hamming_rate |
|---|---|---|
| hapdup_HG002_dual | 0.0418 | 0.24119 |
| shasta_hic_gfase_HG002_standard | 0.11606 | 0.10936 |
| shasta_trio_gfase_HG002_standard | 0.1161 | 0.10727 |
| shasta_hic_gfase_HG002_UL | 0.09252 | 0.07675 |
| shasta_trio_gfase_HG002_UL | 0.09146 | 0.07736 |
| shasta_hic_gfase_HG002_UUL | 0.08471 | 0.07109 |
| shasta_trio_gfase_HG002_UUL | 0.08604 | 0.07304 |
| shasta_hic_gfase_HG002_r10_1fc | 0.05196 | 0.0537 |
| shasta_trio_gfase_HG002_r10_1fc | 0.05061 | 0.04522 |
| shasta_hic_gfase_HG002_r10_ul_4fc | 0.03234 | 0.0241 |
| shasta_trio_gfase_HG002_r10_ul_4fc | 0.03353 | 0.02557 |
| shasta_hic_gfase_HG002_r10_uul_11fc | 0.0336 | 0.0269 |
| shasta_trio_gfase_HG002_r10_uul_11fc | 0.03594 | 0.0285 |
| verkko_gfase_HG002_full_coverage_v1_1 | 0.00639 | 0.00553 |
| verkko_gfase_HG002_production_v1_1 | 0.00621 | 0.00521 |
| verkko_trio_HG002_full_coverage | 0.00846 | 0.00645 |
| verkko_trio_HG002_production | 0.00909 | 0.00639 |
| hifiasm_gfase_HG002_r366 | 0.00643 | 0.0064 |
| hifiasm_hic_HG002_v016 | 0.01105 | 0.01421 |
| hifiasm_trio_HG002_v016 | 0.00822 | 0.00748 |

**Supplementary Table 2:** Evaluation of assemblies using yak Trio eval, comparing existing hybrid and trio assemblers to the HiC phased assemblies presented in this paper.

| **Variant Type** | **Shasta Confident Bed Variants** | **Verkko Confident Bed Variants** | **Shared Shasta Loci** | **Shared Verkko Loci** | **Shasta Only** | **Verkko Only** | **Shasta Indel** | **Shasta Indel Rate** | **Verkko Indel** | **Verkko Indel Rate** | **WhatsHap Covered Variants** | **WhatsHap Switch Rate** | **Whatshap Hamming Rate** |
|---|---|---|---|---|---|---|---|---|---|---|---|---|---|
| 0\|1 | 1282348 | 1397762 | 1257356 | 1262499 | 24985 | 135255 | | | | | | | |
| 1\|0 | 1283631 | 1399130 | 1258035 | 1263396 | 25593 | 135729 | | | | | | | |
| 1\|2 | 42310 | 67573 | 38106 | 32437 | 4204 | 35136 | | | | | | | |
| 2\|1 | 41251 | 67426 | 37137 | 31605 | 4114 | 35820 | | | | | | | |
| Total Hets | 2649540 | 2931891 | 2590634 | 2589937 | 58896 | 341940 | 34364 | 0.01297 | 256474 | 0.0869 | 2,483,238 | 0.000431 | 0.000347 |
| Total Homs | 2046423 | 1749922 | 1665743 | 1666205 | 380680 | 83717 | 342224 | **0.16723** | 30621 | **0.0172** | | | |

**Supplementary Table 3:** Investigation of Dipcall variants for the shasta_trio_gfase_HG002_r10_ul_4fc and verkko_trio_HG002_production assemblies. The assemblies share similar numbers of heterozygous and homozygous variant loci. However, compared to Shasta Dipcall variants the Verkko assembly has more variants, five times more assembly specific heterozygous variants, and seven times as many assembly specific heterozygous Indel variants. While the Shasta assembly has 4.5 times as many assembly specific homozygous variants, and 11 times as many assembly specific homozygous Indel variants compared to Verkko.

We ran WhatsHap Compare with the Verkko assembly as the truth set and Shasta as the query, the last three columns in the table. WhatsHap reports a blockwise hamming rate of 0.00035 for the 2.48 million Verkko heterozygous variants that agree in genotype with heterozygous Shasta Dipcall variants at the same loci.

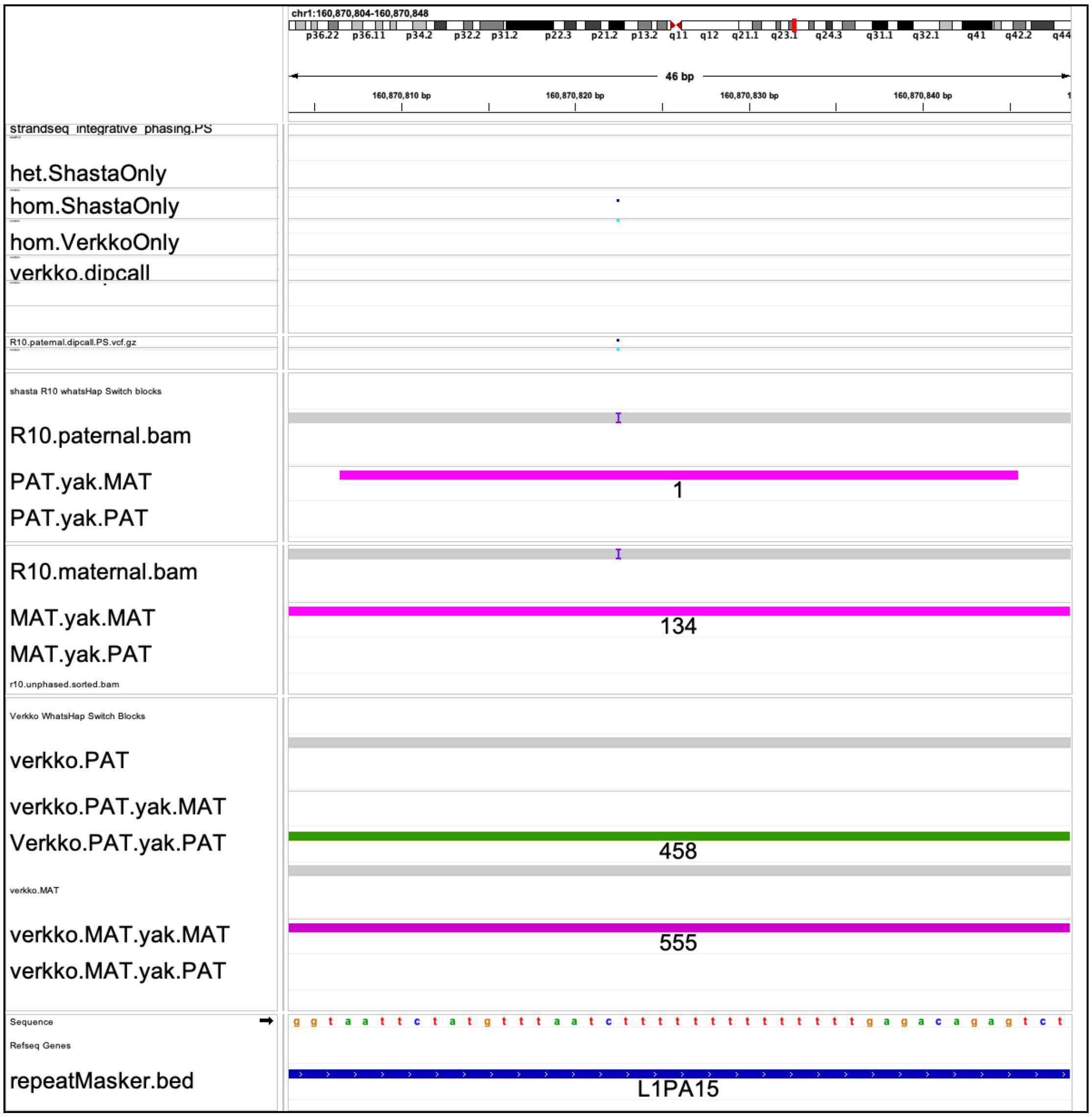


a)

b)

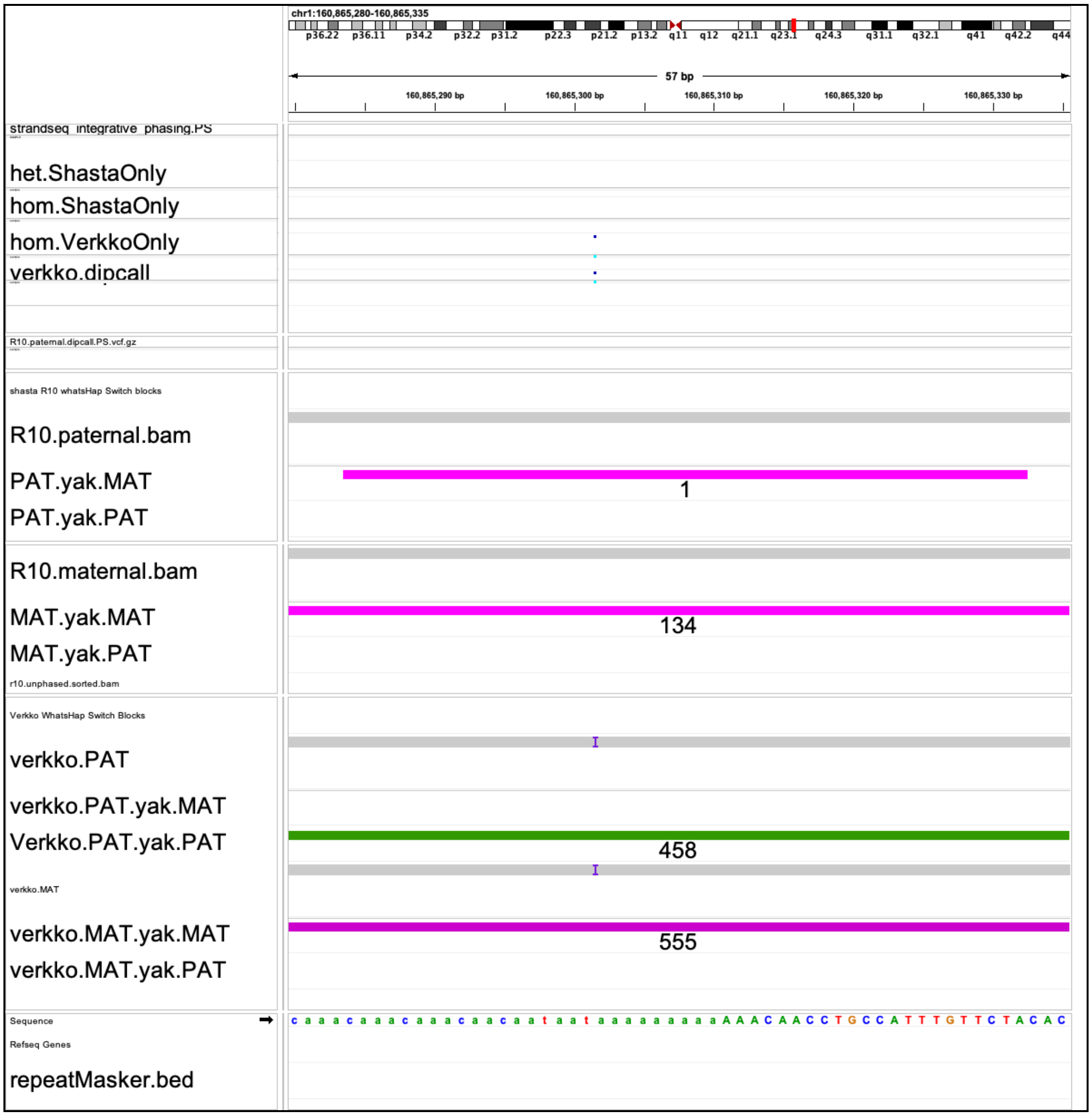
chr1:160,865,280-160,865,335
57 bp
160,865,290 bp
160,865,300 bp
160,865,310 bp
160,865,320 bp
160,865,330 bp
strandseq_integrative_phasing.PS
het.ShastaOnly
hom.ShastaOnly
hom.VerkkoOnly
verkko.dipcall
R10.paternal.dipcall.PS.vcf.gz
shasta R10 whatsHap Switch blocks
R10.paternal.bam
PAT.yak.MAT
1
PAT.yak.PAT
R10.maternal.bam
MAT.yak.MAT
134
MAT.yak.PAT
r10.unphased.sorted.bam
Verkko WhatsHap Switch Blocks
verkko.PAT
verkko.PAT.yak.MAT
Verkko.PAT.yak.PAT
458
verkko.MAT
verkko.MAT.yak.MAT
555
verkko.MAT.yak.PAT
Sequence
Refseq Genes
repeatMasker.bed
c a a a c a a a c a a a c a a c a a t a a t a a a a a a a a a A A A C A A C C T G C C A T T T G T T C T A C A C
c)

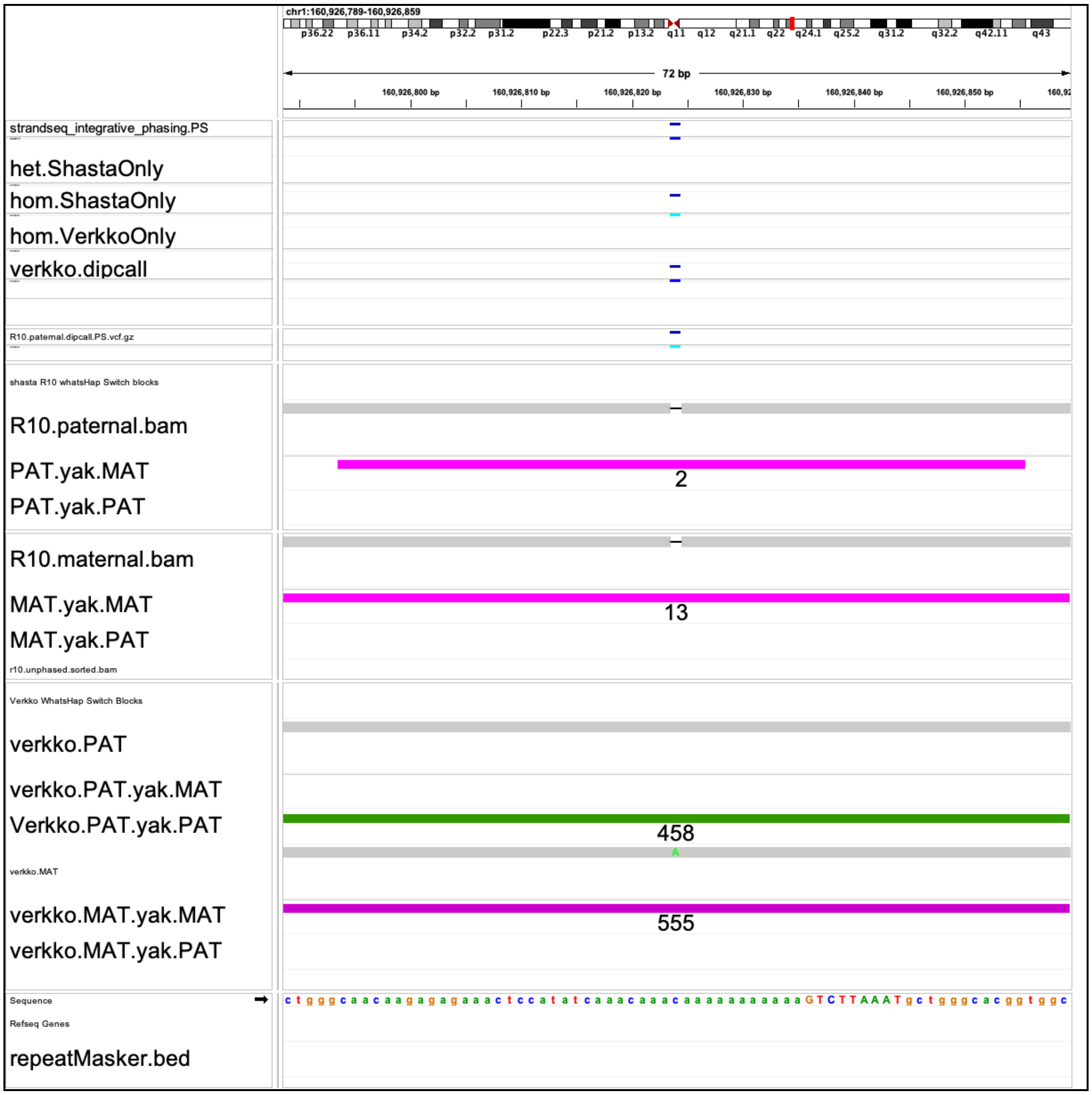
chr1:160,926,789-160,926,859
72 bp
160,926,800 bp
160,926,810 bp
160,926,820 bp
160,926,830 bp
160,926,840 bp
160,926,850 bp
strandseq_integrative_phasing.PS
het.ShastaOnly
hom.ShastaOnly
hom.VerkkoOnly
verkko.dipcall
R10.paternal.dipcall.PS.vcf.gz
shasta R10 whatsHap Switch blocks
R10.paternal.bam
PAT.yak.MAT
PAT.yak.PAT
2
R10.maternal.bam
MAT.yak.MAT
MAT.yak.PAT
13
r10.unphased.sorted.bam
Verkko WhatsHap Switch Blocks
verkko.PAT
verkko.PAT.yak.MAT
Verkko.PAT.yak.PAT
458
verkko.MAT
verkko.MAT.yak.MAT
verkko.MAT.yak.PAT
555
Sequence
Refseq Genes
repeatMasker.bed
ctgggcaacaagagagaaactccatatcaaacaaacaaaaaaaaaaaGTCTTAAATgctgggcacggtggc

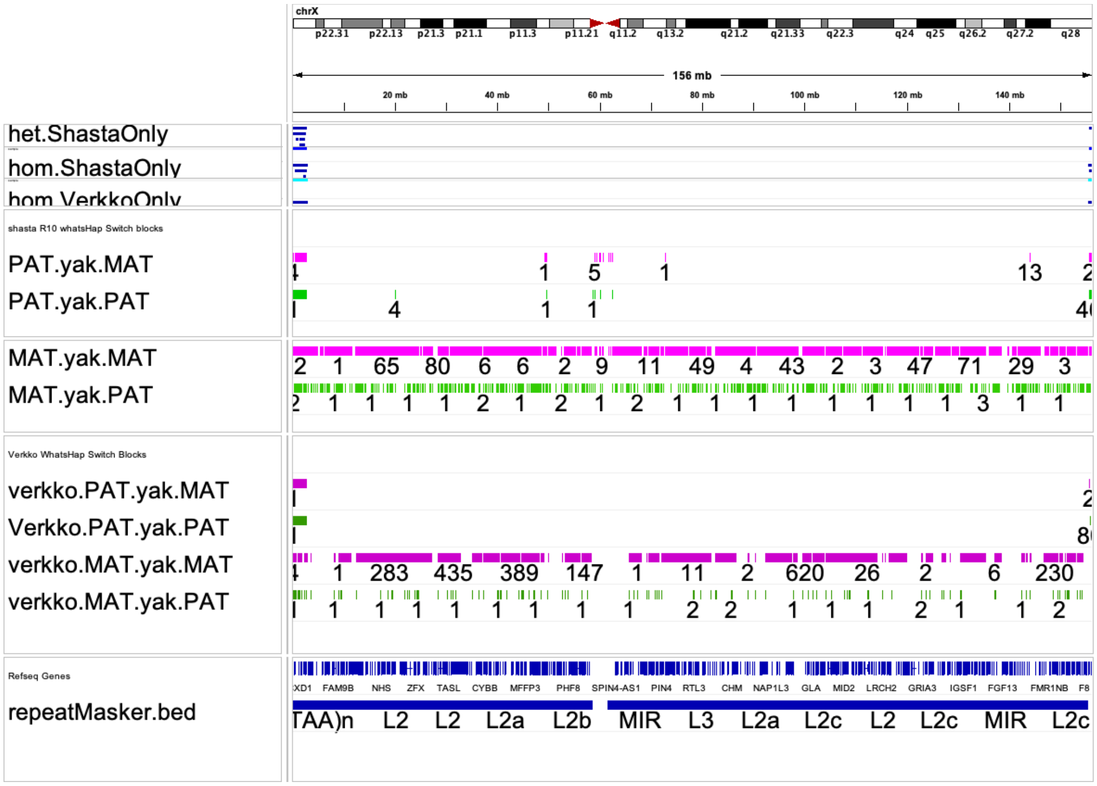


d)

**Supplementary Figure 4:** IGV screenshots of k-mer block locations where homopolymer Indel variants are matching with parental illumina k-mers, as reported by yak on and off target k-mer blocks. Shasta Assembly is shasta_trio_gfase_HG002_r10_ul_4fc, Verkko Assembly is verkko_trio_HG002_production. Magenta bed tracks are maternal yak k-mer blocks labeled with k-mer counts per block. Green bed tracks are paternal k-mer blocks with their respective k-mer counts labeled. a. Shasta specific homozygous “T” insertion in a homopolymer region creates a single maternal k-mer yak switch block in the paternal assembly. b. Verkko homozygous “A” insertion in a homopolymer region creates a single maternal k-mer yak switch block in the paternal assembly. c. A single maternal k-mer yak switch block in the paternal assembly is located at a truth variant. However, the genotypes between assemblies don’t match ( homozygous “CA” deletion in Shasta, C-A SNP in Verkko ) and as such wouldn’t be assessed by WhatsHap. d. Chromosome X illustrates a thicker lawn of off target green paternal k-mer matches in the Shasta yak blocks compared to the Verkko paternal yak k-mer blocks.
